# Multi-scale phase separation by explosive percolation with single chromatin loop resolution

**DOI:** 10.1101/2022.04.28.489670

**Authors:** Kaustav Sengupta, Michał Denkiewicz, Mateusz Chiliński, Teresa Szczepińska, Ayatullah Faruk Mollah, Sevastianos Korsak, Raissa D’Souza, Yijun Ruan, Dariusz Plewczynski

## Abstract

The 2m-long human DNA is tightly intertwined into the cell nucleus of the size of 10μm. The DNA packing is explained by folding of chromatin fiber. This folding leads to the formation of such hierarchical structures as: chromosomal territories, compartments; densely packed genomic regions known as Chromatin Contact Domains (CCDs), and loops. We propose models of dynamical genome folding into hierarchical components in human lymphoblastoid, stem cell, and fibroblast cell lines. Our models are based on explosive percolation theory. The chromosomes are modeled as graphs where CTCF chromatin loops are represented as edges. The folding trajectory is simulated by gradually introducing loops to the graph following various edge addition strategies that are based on topological network properties, chromatin loop frequencies, compartmentalization, or epigenomic features. Finally, we propose the genome folding model - a biophysical pseudo-time process guided by a single scalar order parameter. The parameter is calculated by Linear Discriminant Analysis. We simulate the loop formation by using Loop Extrusion Model (LEM) while adding them to the system. The chromatin phase separation, where fiber folds into topological domains and compartments, is observed when the critical number of contacts is reached. We also observe that 80% of the loops are needed for chromatin fiber to condense in 3D space, and this is constant through various cell lines. Overall, our *in-silico* model integrates the high-throughput 3D genome interaction experimental data with the novel theoretical concept of phase separation, which allows us to model event-based time dynamics of chromatin loop formation and folding trajectories.

## 1. Introduction

The human genome, during interphase, is hierarchically packed in the nucleus with structures at different scales, organizing DNA into functional components (Tang *et al*., 2015; Kempfer and Pombo, 2020; Zheng and Xie, 2019; Pombo and Dillon, 2015) (Fig. 1A). Chromatin at the highest scale is separated into two major compartments: compartment A, where the chromatin has a wider spacing between the nucleosomes and is more transcriptionally active, enriched by histone modifications, such as H3K4me3, H3K27ac, H4K16ac; and compartment B, where the fiber is more tightly packed and transcription is limited (Lieberman-Aiden *et al*., 2009; Rao *et al*., 2014; Kumar *et al*., 2019). Microscopy studies show that the compartments are generally spatially separated in the cell nucleus (Shiels *et al*., 2007). The next scale in the hierarchy is the existence of the chromosomes, which occupy distinct territories in the nucleus of the cell (Dorier and Stasiak, 2009; Meaburn and Misteli, 2007) and only partially overlap (Branco and Pombo, 2006). The chromatin is further divided into domains with a relatively large number of internal contacts, called Topologically Associating Domains (TADs) (Nora *et al*., 2012; Dixon *et al*., 2012; Dekker, 2016) Chromatin Contact Domains (CCDs) (Tang *et al*., 2015). These contacts include chromatin loops mediated by factors like CCCT-binding factor (CTCF) and cohesin, or enhancer-promoter contacts (Fig. 1B). Compartmentalization and condensation, while being general principles of genome organization, simultaneously play a major role in determining the functionality of the cell. The biophysical process behind the compartmentalization is proposed to be phase separation (Gibson *et al*., 2019; Narlikar, 2020) (Fig. 1C). Phase separation in biology is a biophysical process of spontaneous separation into a dense and dilute phase of a well-mixed solution of biomolecules (proteins, nucleic acids, chromatin fiber). This phenomenon gives rise to diverse non-membrane-bound nuclear, cytoplasmic, and extracellular compartments (Hyman *et al*., 2014; Noda *et al*., 2020). A phase separation model of microcompartment formation was proposed for both transcriptionally active (Hnisz *et al*., 2017) and heterochromatin regions (Strom *et al*., 2017).

**Figure 1:**
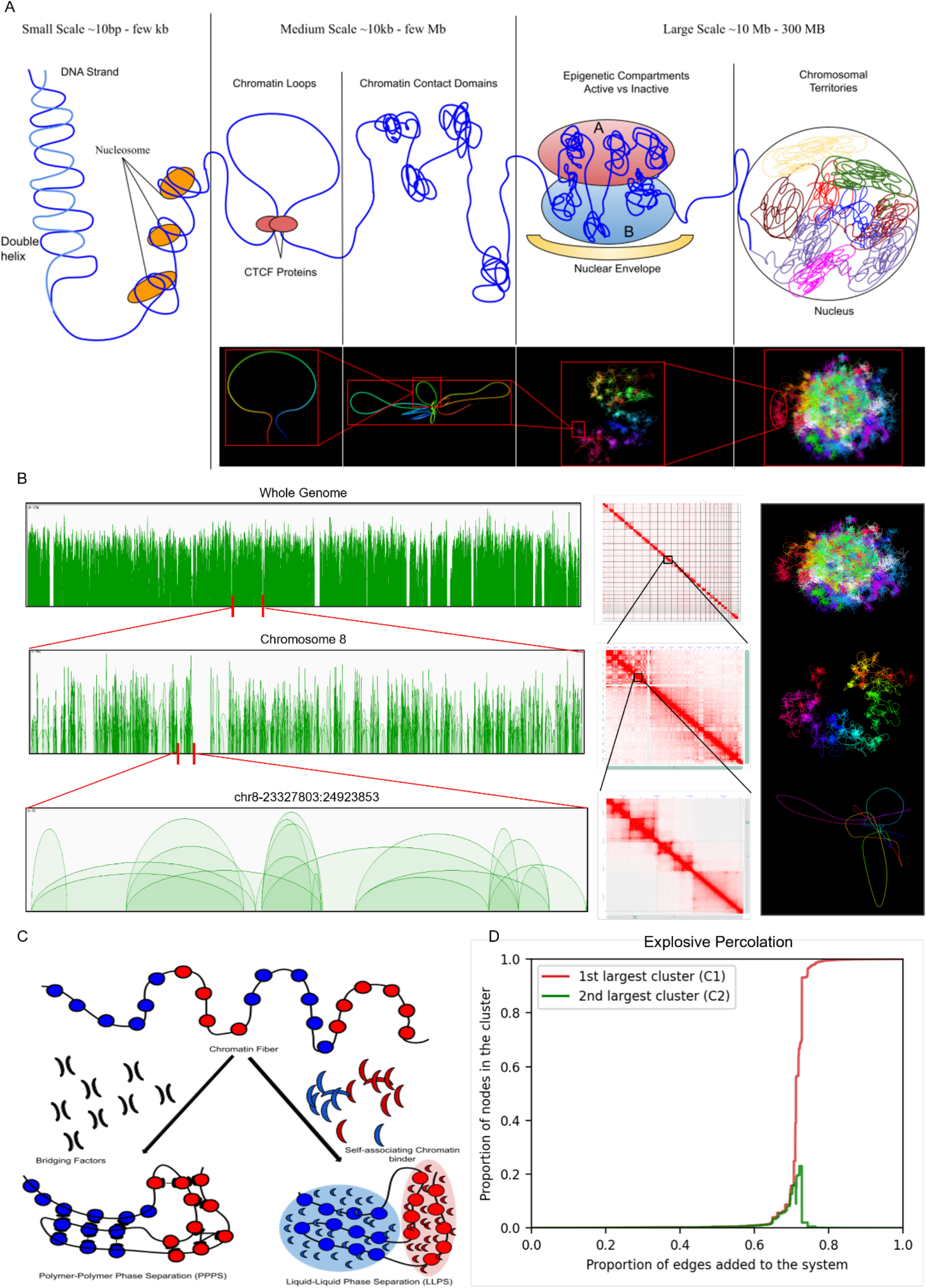
(A-B) Shows the schematic diagram, arc diagrams, heatmaps and 3D models of multiscale representation of the genome. (C) Shows the schematic representation of Polymer Polymer Phase Separation (PPPS) and Liquid Liquid Phase Separation (LLPS). (D) Shows a typical evolution of cluster size for adding edges to a graph for first and second largest cluster using the Frequency Model (FM) of edge addition.

During their life cycle, cells undergo a differentiation into specialized cell types, leading to a structural reorganization of the chromatin (Zheng and Xie, 2019; Sun *et al*., 2018; Slack, 2007). In recent years, a number of methods have been proposed to unravel the genome-wide chromatin folding patterns (Tang *et al*., 2015; Lieberman-Aiden *et al*., 2009; Tavares-Cadete *et al*., 2020; Zheng *et al*., 2019). The most commonly-used techniques are based on the Chromatin Conformation Capture (3C) method and allow the identification of chromatin contacts genome-wide (Han *et al*., 2018). The two major examples include Hi-C (Lieberman-Aiden *et al*., 2009; Rao *et al*., 2014) and Chromatin Interaction Analysis by Paired-End Tag sequencing (ChIA-PET) (Tang *et al*., 2015; Li *et al*., 2010). While Hi-C captures unspecific contacts and works in lower resolutions (up to 1kbps), ChIA-PET concentrates on detecting interactions mediated by specific proteins, allowing detections of loops in very high resolution (of up to 100bps). While Hi-C provides information about strength of connection for all-vs-all genomic regions, ChIA-PET shows only the specific interactions mediated by the target protein.

Understanding the folding of DNA into hierarchical components from the loops or contacts detected by various methods is an open research area. Various mathematical models have been proposed over time to predict the folding of DNA inside the nucleus. Emmett et al. created the first method based on a topological data analysis to understand the structure of Hi-C data (Emmett *et al*., 2016). However, that study was restricted to one chromosome at 1Mb resolution due to computational constraints. Recently Carriere and Rabadan used topological data analysis to explore the similarity between two single-cell Hi-C maps (Carrière and Rabadán, 2020). Unfortunately, neither of the methods proposes the existence of a general mechanism on how the hierarchical structures are formed while folding of the whole genome nor do they consider the dynamic nature of chromatin loop formation.

Percolation is the process of adding connections to a network, which leads to a sudden emergence of connectedness in a significant part of the network; this sudden emergence of connectedness is called the percolation phase transition in graphs (M. Sahimi, *Applications of Percolation Theory*, Taylor & Francis, London, 1994., A. A. Saberi, Phys. Rep. 578 (2015)). Phase transitions can be classified as either first or second order, based on their fundamental characteristics. The first-order phase transition is discontinuous, abrupt, and can have “explosive” behavior (Fig. 1D), whereas the second-order phase transition is continuous and smooth, with the sizes of the clusters generally following a power-law distribution. Percolation theory has been used to study a wide range of real-world network systems such as social networks (Chen *et al*., 2007) or signal transmission between bacteria (Larkin *et al*., 2018). In particular, explosive percolation emerged recently as a powerful approach which shows how low-level random processes can strongly impact the phase transition in the large-scale system (Achlioptas *et al*., 2009; D’Souza and Nagler, 2015; D’Souza *et al*., 2019).

The main biological mechanism that is responsible for the formation of TADs is considered to be the Loop Extrusion process (LE). The loop extrusion is mediated by a loop extrusion factor, which usually is a ring-like Structure Maintenance of Chromatin complex (SMC), which extrudes a loop bidirectionally with a speed of ∼0.5-1.0 kb/second (Hansen, 2020). The motion of an SMC complex stops when it meets a boundary element (BE). Cohesin or condensin usually play the role of loop extrusion factors, and they can be blocked by another loop extrusion factor (Banigan and Mirny, 2019) or a boundary element like a CTCF protein (Xiao *et al*., 2011). Interestingly, CTCF proteins are connected with 11-Zinc finger motif sequences of nucleotides, and, therefore, we can define an orientation according to these sequences. CTCF motifs are enriched in the boundaries of TADs with (usually) convergent orientation, which means that loop extrusion is usually mediated by a SMC complex that extrudes a loop and is bounded by two CTCF motifs in convergent direction (Lazniewski *et al*., 2019). It may also happen that cohesin will meet a CTCF in an opposite orientation than the direction of the movement of cohesin. In this case, cohesin slows down when it meets it and passes through it. This is the reason why sometimes dots exist in the boundaries of TADs. Loop Extrusion is connected to TADs, but the relation is not clear, since TADs express a superposition of many biological processes (Hansen, 2020).

Some interesting questions regarding the chromatin folding are: what is the exact path or trajectory of the folding, what are the minimum percentages of loops required for the chromatin to fold correctly, and what is the interplay of loop formation and folding on various scales of hierarchical structures of chromatin. In this study, we introduce a graph-based method using percolation theory and dynamics of loop extrusion to simulate the formation of chromatin loops and the folding of chromatin fiber, by adding loops to the system over high-resolution data at several levels of the hierarchical structure.

Furthermore, we propose various theoretical models of percolation over graphs created from 3D chromatin interactions. The chromosomes are modeled as graphs where CTCF chromatin loops are represented as edges. The folding trajectory is simulated by gradually introducing loops to the graph following various edge-addition strategies that are based on topological network properties, chromatin loop frequencies, compartmentalization, or epigenomic features. For each instance of loop selected by percolation, we use loop extrusion to extrude the loop, thus adding the dynamics of loop forming to the mix. This gives us the interplay between the macro scale (compartments) and the micro scale (TADs) folding of chromatin. In our models, the percolation gives the macro scale folding to compartments and loop extrusion gives us the micro folding to TADs. We compare the models obtained from various edge addition techniques of percolation to find the most suitable model to show the trajectory followed by chromatin folding into the higher structures such as compartments and TADs. Overall, in the proposed work we show that the trajectory of chromatin folding in the nucleus and the formation of hierarchical structures like compartments and CCDs can be inferred in pseudo-time, using the concepts of explosive percolation theory, phase separation, and loop extrusion over 3D chromatin interaction graphs. We also show that at least 80% of loops are required for correct folding in 3D space and this threshold remains constant across different cell lines.

## Methods

### 3D structure data

In the present study, we focus on ChIA-PET assay obtained from (Tang *et al*., 2015), mapping interactions mediated by CTCF to hg19 reference genome for human lymphoblastoid cell line GM12878. The number of times each interaction has been observed in the experiment (loop frequency) is called the PET count - the higher the PET count, the more reliable the interaction. In this study we only include intrachromosomal interactions with PET count equal to or above 2. For every chromosome we create a network where each ChIA-PET interaction is represented as an edge between two ends of a loop (anchors). If a group of anchors has overlapping coordinates, we merge them into a single node in the network. Moreover, we connect linearly consecutive anchors with an edge called Linear Edge (Fig. 2A). We focus on chromosomes 1 to 22, excluding the X and Y chromosomes due to the limitations of the interaction data, thus obtaining 22 separate networks - one for each of the autosomes. We also use chromatin features from ENCODE to guide the percolation in feature-based models (see Supplementary Text).

**Figure 2:**
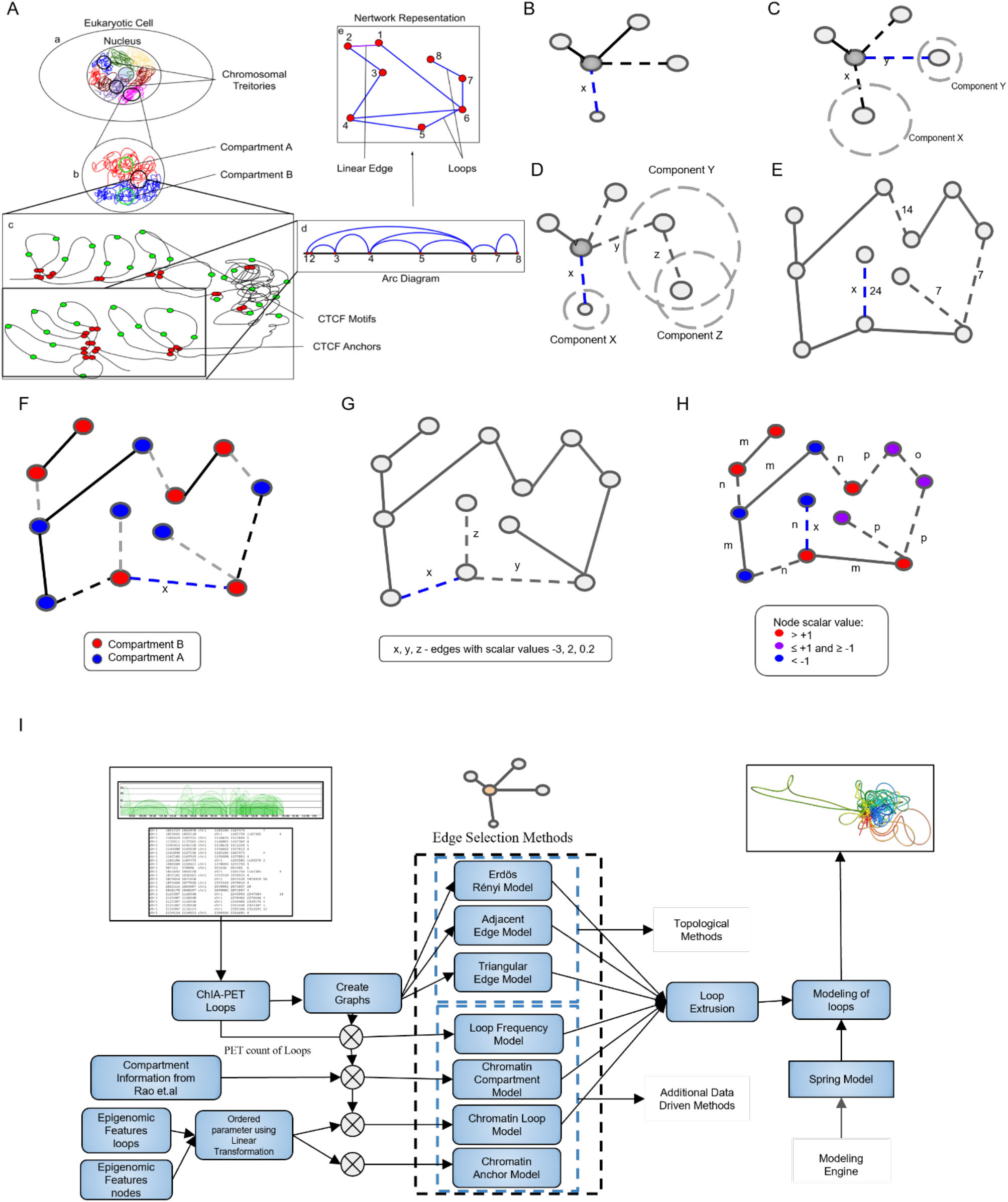
(A) Chromatin can be represented as a graph. ChIA-PET anchors are represented as nodes (red) and loops as edges (blue). If two adjacent anchors have no loop between them, we put a linear edge representing the chromatin fiber (magenta). (B-H) Various edge addition methods. Dashed lines are ChIA-PET loops not added to the graph yet. Blue, dashed line indicates chosen edge: (B) **Erdös-Rényi (ER) model**. In ER an edge is selected uniformly at random and added to the graph (e.g. edge x). (C) **Adjacent Edge (AE) model:** In AE we select at random two edges connected to a single node v and add the edge connected to the smaller component (y is chosen, as component Y is smaller). (D) **Triple Edge (TE) model:** In TE we randomly select three candidate edges: x, y and z, where x and y share a common node v and z is connected to either x or y and we select the edge connected to the smallest component (edge x, because X is the smallest component). (E) **Loop Frequency (LF) model:** In LF the process of edge addition is done according to the frequency of the ChIA-PET loops. The edges with the highest frequency are added first (edge x). If the edges have equal frequencies, one is chosen from them randomly. (F) **Chromatin Compartment model (CC):** We classify the nodes into compartment A or compartment B. We add the intra-compartmental loops (coloured black) first, according to their frequency. Next, we add the inter-compartmental (colored grey) loops also following the frequency order. (G) **Chromatin Loop model (CL):** Here we use linear regression to calculate a single scalar value that classifies loops into compartments based on chromatin features. We add edges according to the absolute value of the scalar from the highest to the lowest (let’s assume if x has a scalar value -3, y has a scalar value 2 and z has a scalar value 0.2 then we add in order x, y, z to the system). (H) **Chromatin Anchor Model (CA)**: In the first step we look at the scalar value calculated for anchors and divide the nodes into three sets, based on whether the value is greater than one (red, set 1), less than minus one (blue, set 2), and the rest (purple, set 3). We then add edges in several stages depending on the sets to which their ends belong: both in set 1 (type m), both in set 2 (also type m), between set 1 and set 2 (type n), both in set 3 (type o), and finally between set 3 and sets 1 and 2 (type p). At each stage we find the edge with the smallest possible difference in the anchor scalar values and add that edge to the system. If there are edges with the same difference of anchor scalar value, then we add the edge with the highest absolute loop scalar value. If this value turns out to be equal for both edges, then we choose the edge to add randomly. (I) Chromatin Folding Simulation using Percolation with Loop Extrusion Model in one modeling approach.

Furthermore, to show that the method is constant across datasets of features and different cell lines, we used 4DN cell lines (GM12878, H1ESC, and HFFc6; all mapped to hg38) and Avocado (Schreiber *et al*., 2020) imputed epigenetic marks for further validation. In this study, we used 69 epigenetic marks (see Supplementary Table 2) obtained for GM12878, 84 features for H1 cell lines, and 82 epigenetic marks for HFF cell lines using Avocado software.

### Percolation mechanism and the critical point

A *connected component* in a network is a set of interconnected nodes, separated from the rest of the network. More specifically, it is defined as the maximum set of nodes, such that for any two nodes *u* and *v* in this set, a path exists from *u* to *v*. We refer to connected components as *clusters*.

As edges are gradually added to the system, the clusters merge together and grow in size. To quantify the changes that occur in the system during this process we keep track of three parameters: the size of the largest and second-largest clusters in the network (*C*_*1*_ and *C*_*2*_ respectively), and the second moment of cluster sizes (*m*_*2*_). The second moment can be understood as the expected size of the connected component that contains a chosen vertex, and can be defined as follows:

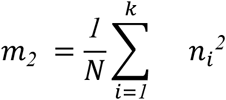

where *n*_*i*_ is the number of nodes in the i-th clusters, k is the number of distinct clusters, and *N* is the total number of nodes in the network. The parameters are most conveniently expressed as normalized to the range *0-1*, by dividing by *N*.

the percolation process starts with a fully disconnected network, without any edges, in which each node forms a distinct, connected component, and all three parameters have minimum values of almost *0*. Then, edges are gradually added to the system, according to a chosen method, causing the clusters to merge together and grow in size. As this happens, *C*_*1*_ and m_2_ grow accordingly, finally reaching their maximum value of *1* when the network becomes fully connected, i.e. all the nodes are in a single giant cluster. The trajectory of *C*_*2*_ is more complicated, as it initially rises, and then needs to drop to *0* as the two largest clusters encounter each other and merge. Percolation theory characterizes types of percolation on the basis of the trajectories of these parameters (eg. (D’Souza and Mitzenmacher, 2010)). In particular, in case of *explosive percolation*, after the critical number of edges is added, a dramatic change in the parameter trajectories occurs: both *C*_*1*_ and *m*_*2*_ sharply increase, and *C*_*2*_ begins to drop. This corresponds to the network rapidly transitioning from being mostly disconnected islands, to a state with a single dominating cluster.

### Multi-scale Monte Carlo Feature Selection (MCFS)

We used the Monte Carlo Feature Selection (MCFS) algorithm (Dramiński and Koronacki, 2018) for the importance analysis of the epigenetic features. First, we took Avocado imputations (Schreiber *et al*., 2020) for various epigenetic marks (84 in total for H1, 69 features for GM12878, and 82 for HFF cell lines - see Supplementary Table 2) and create an epitensor of 1kbps resolution. Based on that epitensor, we added the class depending on the compartment. The MCFS algorithm then used the epitensor for the establishment of the relevance index (RI) of each of the features. The epitensor was divided into *s* subsets of m epigenetic features, and each subset created *t* splits of the data into training/testing sets and used that for training a decision tree. Based on all the information from the splits in totally-*st* trees, the relevance of a particular feature is calculated. Such information is then used for creating the importance ranking, and dependency graphs which show the dependencies of the epigenetic marks on each other. The bigger the node, the more important the feature in a given classification task, and the bigger the arrow pointing from it, and the more this particular node (epigenetic mark) influences other nodes (another epigenetic mark).

### Scalar value of compartmentalization

We computed the scalar value (Supplementary Text) for loops and anchors separately. We collected a range of genomic data that characterize chromatin state and chromatin binding proteins in GM12878 cells and constructed 428 features for GM12878 hg19 by calculation of the fraction of the loop/anchor covered with the peak signal, the mean, or the sum of the peak signal along the whole loop/anchor or the mean of the whole signal along the loop without peak identification, counted as separate features. For validation with other cell lines, we used 69 features for GM12878 hg38, 84 features for H1ESC hg38 cell lines, and 82 features for HFFc6 hg38 cell lines and selected the most important features using MCFS. Over these sets of features, we compute the scalar value by constructing a hyperplane *H* that optimally separates loops/anchors of compartments A and B, using Fisher’s linear discriminant analysis (Fisher, 1936). Then, we consider the orthogonal distance between *H* and a loop/anchor, represented as point x in n-dimensional space. We can denote this distance value as *d = w*^*T*^*x*, where *w* is the vector normal to *H*. For all points *x* that belong to the hyperplane, the value of *d = 0*. Based on the value of *d* we can label the data into three sets, first of loops/nodes, second of loops/nodes, and the last of intercompartmental loops/nodes. Moreover, the higher the absolute value of *d*, the higher the probability of it belonging to one of the compartments. Based on the scalar parameter value we classify loops and nodes into set 1, having scalar parameter value greater than 1, set 2 - if the value is less than -1, and the remaining ones to set 3. set 1 and set 2 contain loops/nodes from a specific compartment and set 3 contains the loops/nodes which are classified as missing. We use values greater than 1 or less than -1 because this gives us best accuracy.

### Topological models

At each time point a model selects a single edge to be added to the graph. An edge can be added between nodes *u* and *v* if it is present in the original network, i.e. there is a ChIA-PET loop between them or they are linearly consecutive in the sequence. **Erdös-Rényi (ER) model (Fig. 2B):** The edge is selected at random with equal probability. **Adjacent Edge (AE) model (Fig. 2C):** Two edges sharing a common vertex *u* are initially selected uniformly at random. The addition of either would connect *u* to a different connected component; We choose the one that would connect *u* to the smaller of the two components and add it to the graph. If the components have the same size (or are, in fact the same component) one of the two edges is chosen randomly. **Triple Edge (TE) model (Fig. 2D):** Since our graphs are sparse (edge-to-node ratio ranges from 1.51 for chromosome 13 to 1.55 for chromosome 19, see also Supplementary Figures 1 to 22), we modify the triangle rule, originally described in (D’Souza and Mitzenmacher, 2010). In this method we first select three candidate edges. The first two edges, *(u, v)* and *(u, w)*, sharing a vertex *u*, are selected uniformly at random, as in the AE method. Then, the third edge is selected at random in such a way that one of its vertices is *v* or *w*. Finally, out of the three candidate edges, the one connecting to the smallest component is added to the graph. Our experiment is more inclined with the models stated in (D’Souza and Mitzenmacher, 2010) and more biologically plausible rather than approaching more complex random network models. Thus the models detailed in (D’Souza and Mitzenmacher, 2010), serve as a baseline for the following proposed models driven by additional data.

### Loop Frequency model

While the aforementioned models use interactions to form graphs, they do not consider the strength of the interaction in selecting the new edge. To fully utilize the available information, we proposed the Loop Frequency model (LF; Fig. 2E), in which the edges are added according to the loop frequency (indicating the strength of the interaction) in ChIA-PET data. This way the most stable or the most populated loops are added first.

### Chromatin Compartment model

To study and reproduce the biology behind compartmentalization, we extend the physical idea of edge addition of percolation theory using the biophysical concept of block polymer. In the Chromatin Compartment model (CC; Fig. 2F), the polymer is represented by node clusters that belong to the same compartment: A or B. We classify the loops into two types: inter-compartmental and intra-compartmental and add intra-compartmental loops into the graph first. Among the intra-compartmental set of loops, we first select those with the highest frequency. After all intra-compartmental loops are added, we add the inter-compartmental loops following the loop frequency as well. This model captures the genomic phase condensation and separation into compartments.

### Chromatin Loop model

In the Chromatin Loop model (CL, Fig. 2G) we used the scalar value of loops (see scalar value of compartmentalization section and Supplementary Material) to guide the percolation trajectory. We started from the edges with the largest absolute value of the scalar value, which means edges strongly associated with either compartment A or B. Then, we gradually added edges with lower scalar values. Thus, as we approach the value of *0*, we include more loops that are not clearly associated with either compartment. Therefore, in this model we obtain a separate phase for each compartment. During our experiments we found that this model is sensitive to the topology of the graphs generated for each chromosome and we observe high inter-chromosomal variations. Therefore, we extend the results with the Chromatin Anchor model (discussed in the next section) with a scalar parameter for both anchors and loops to capture the biology behind the chromatin folding more accurately.

### Chromatin Anchor model

In the next step, we introduce the Chromatin Anchor model of percolation (CA; Fig. 2H). In this model, we used the scalar values of both nodes (anchors) and edges (loops) to guide the percolation. We took a group of nodes with similar scalar values from either *set 1* or *set 2*. If the difference between scalar parameters of the anchors is equal for two loops, we select the one with the highest absolute value of scalar value. Having added all the loops between nodes from *set 1* and *set 2*, we add loops associated with nodes from *set 3*. For *chromosome 8* we have *32142* nodes and *17060* loop edges and *31805* linear edges of the *32142* nodes, *10669* fall in *set-1, 15202* nodes fall in *set-2* and *12692* in the *set-3*. In CA, we guide the percolation using only the information about interactions, without factoring in the strength or the compartmentalization. We can consider the CC model as the ground truth, as it is purely based on the compartments detected in Hi-C, and we can observe it using our predicted scalar parameter, and we can reconstruct the same trajectory as the CC model.

### Quantifying Percolation

As the primary measure of percolation, we use the size of the largest cluster (*C*_*1*_) normalized to the range *0 - 1* (i.e., the proportion of nodes in the largest connected component). For each percolation model, we construct *1000* trajectories on a given graph (chromosome) and construct an averaged trajectory by taking the mean value in each time point. In all subsequent analyses, we use these averaged trajectories. To quantify the rapidness of percolation we use two complementary approaches. The first approach, meant to be robust, is based on the observation that once the largest cluster contains a significant proportion of all nodes, percolation enters an explosive stage, and later slows down just before most of the nodes are joined. We define the onset of the explosive stage as the point at which the largest connected component size reaches *10%* of the total number of nodes. In the same manner, offset is defined as the point where *90%* of nodes are connected. We also define the critical region width as the difference between the offset and onset. In the second approach we quantify the rapidness of growth of the largest cluster directly, by fitting a univariate linear regression model to the points in the trajectory within the critical region (between the onset and offset). From this regression model we obtain the slope value (regression coefficient); The larger the slope, the more rapid percolation. Moreover, we use the bias parameter, which is the point of the regression line crossing the *C*_*1*_ value of *0*.*5* (Fig. 1D). Finally, in order to validate our detection of the critical region, we track the relative size of the second-largest cluster (*C*_*2*_), and the normalized second raw moment of cluster sizes (*m*_*2*_); both measures should exhibit an increase in the critical region, with *C*_*2*_ dropping to *0* after the initial increase.

### Phase Separation

The compartmentalization can be defined as a phenomenon of microphase separation of polymer blocks called polymer-polymer phase separation (PPPS) (Hildebrand and Dekker, 2020). We define the phase separation in our models as a separated formation of two clusters in the percolation trajectory, following this definition of PPPS (Dekker, 2016; Boeynaems *et al*., 2018). The fundamental physical model behind this process is a polymer made up of small blocks of monomers that belong to different separately-phasing compartments (Dekker, 2016; Dekker and Mirny, 2016; De Gennes and Gennes, 1979; Leibler, 1980; Matsen and Schick, 1994). The polymer can be viewed as a block polymer, and it can fold independently, i.e. the blocks have significantly low inter-block interaction. In our graphs we classify the nodes/edges into two or more classes depending on the compartment information of the scalar value calculated from a set of genomic features. These clusters tend to have higher intra-cluster edges than inter-cluster edges (Hildebrand and Dekker, 2020).

To replicate the phase separation phenomenon in our proposed folding trajectories, the intra-cluster edges are added to the systems at the first step followed by the inter-cluster edges.

### 3D chromatin models

The 3D chromatin models are built using the Spring Model (SM) software (Kadlof *et al*., 2020). Spring Model uses the mechanism of molecular mechanics and is implemented using the OpenMM - python framework for molecular dynamics simulations. The modeling engine uses the beads-on-chain representation by representing polymers as a set of points in 3D space. Each bead represents an equal amount of polymer in the resolution specified by the user. For whole chromatin models in our work, we use a resolution of *50kbp*, i.e., one single bead represents 50,000 base pairs. The beads which are far apart in the chain are connected using springs (harmonic bonds) if there is an interaction between them in the graph. Then SM simulation performs energy minimization over the forces determined by the springs and chain properties (e.g., rigidity) to determine the 3D shape of the molecule with the set of contacts described by the graphs. We start the modeling with an initial circular 3D structure - without any loops (springs). Then, for each step of the percolation process, we introduce an edge (as added in the percolation model) as a spring to SM, thus bringing the ends of the contact close in 3D space and creating a loop.

### Loop Extrusion Model (LEM)

In Loop Extrusion Model we introduce the idea of dynamic loop formation by single side loop extrusion instead of static loop addition to the system after we select the loop by one of the above-described percolation models. To model the loop extrusion, we create synthetic LEM data which are based on the idea that while the cohesin complex moves through the DNA fiber, it continuously extrudes the loop. We first identify the CTCF motif by the DNA sequence search using matrices from Jasper [44]. Then the motifs are filtered using locations of CTCF ChIP-seq peaks obtained from ENCODE(ENCSR000DZN) [36]. If in a loop at least one CTCF motif is present, we synthetically extrude this loop. In This model, a stopping point of cohesin appears if one of the two conditions are satisfied: (i) cohesin reaches a stopping point marked by the anchors, or (ii) cohesin encounters two CTCF motifs with a convergent orientation. If the orientation of the CTCF motif is divergent, the cohesin slows down. We apply LEM to percolation models at the CCD level due to computational limitations (see Supplementary Figure and Supplementary Video).

## 2. Results

In the proposed method the structure of each chromosome in the human genome is presented as a graph derived from the ChIA-PET experimental data, with edges representing CTCF chromatin loops, and nodes representing the anchors of those loops (Halder *et al*., 2020). This representation of the chromatin spatial structure allows us to study its behavior using a variety of established graph theory methods. Therefore, using high-throughput structural data, we model chromatin hierarchically, starting at the loops, through CCDs, up to compartments (Fig. 2A).

Using these graphs, we simulate the folding of the entire chromosomes with an abstract event-based timeline, where the formation of a single chromatin loop constitutes an event. The order of loop formation follows from various probabilistic models. By studying the percolation of the genomic network, we are exploring how the genome folds when loops are created. we start with an empty network containing only nodes, then keep on adding edges using various strategies, simulating how the chromatin folding process can continue (Fig. 2B-H). Before the phase transition point in percolation, the chromosomes remain unfolded, and after phase transition they are folded in 3D space. We use three standard edge addition methods which operate solely based on network topology: Erdös-Rényi, Adjacent Edge, and a slight modification of Triangular Edge, which we call Triple Edge (D’Souza and Mitzenmacher, 2010; D’Souza and Nagler, 2015; Gilbert, 1959). We then propose models driven by additional experimental data, namely, frequencies of ChIA-PET interactions, information about compartments from Hi-C data, and a scalar parameter that is calculated using linear regression over chromatin features at anchors and loops level. Chromatin features include expression, histone modification, and transcription factor binding. We also find the critical point or: critical percentage of loops which results in complete folding of the chromatin fibers in 3D space.

### Effectiveness of the Scalar Parameter

Our proposed percolation scheme is further guided using biological insights by incorporating genomic features. We have collected a range of genomic data that characterize chromatin state and chromatin binding proteins in GM12878 cells. These are mostly ChIP-seq data of histone modifications (H3K27ac, H3K27me3, H3K36me3, H3K4me1, H3K4me2, H3K4me3, H3K79me2, H3K9ac, H3K9me3, H4K20me1) and transcription factors from ENCODE project (Dunham *et al*., 2012). Additionally, we have collected data about DNA methylation, open chromatin state (DNAse-seq, FAIRE-seq, and ATAC-seq), RNAs (RNA-seq), and nascent RNAs (GRO-seq, Bru-seq). Moreover, we have used ENCODE-combined data: open chromatin synthesis and genome regulatory segmentation (ChromHMM, Segway methods, and compilation of both). We have collected the GC percentage information as well. Each loop is assigned one number for each feature. It is a fraction of the loop covered with the peak signal, the mean, or the sum of the peak signal along the whole loop, or the mean of the whole signal along the loop without peak identification. Therefore, to each genomic dataset, we have assigned several features. Then we use the features to obtain a scalar value, which classifies loops into compartments, to guide the percolation. We have classified loops into compartments by means of linear discriminant analysis (LDA), a supervised machine learning method. To validate the classification from LDA, we compared them with the ground truth of compartments detected using Hi-C data from (Rao *et al*., 2014). AUC score for compartment prediction by the LDA model used to obtain the scalar value is *0*.*95*, which confirms good prediction quality. We checked the correlation of the scalar value calculated for loops with a set of active (DNAse-seq, H3K36me3, H3K27ac, H3K4me2, H3K9ac) and inactive chromatin marks (H3K27me3). H3K36me3 signal is stronger when the scalar value is moving towards a positive value (*slope of linear regression = 0*.*59, spearman correlation = 0*.*70*) (Supplementary Figure). We have calculated the scalar value for anchors in the same way as we have for loops. Classification of anchors into compartments by scalar value is also effective (*AUC score equal to 0*.*89* when compared to the ground truth from (Rao *et al*., 2014)). We have checked the correlation of the scalar value calculated for loops with a set of active (DNAse-seq, H3K36me3, H3K27ac, H3K4me2, H3K9ac) and inactive chromatin marks (H3K27me3). H3K36me3 signal is stronger when the scalar value is moving towards a positive value (*slope of linear regression = 0*.*54, spearman correlation = 0*.*70*). This confirms that the positive value of the scalar reflects the active compartment (Supplementary Fig. 24G and 24H). We observe similar trends in the plots with other active methylation marks as well. In summary, the scalar value efficiently captures the idea of compartmentalization. The findings are also consistent with (Lieberman-Aiden *et al*., 2009).

### Chromatin Compartment, Loop Frequency, Chromatin Loop, and Chromatin Anchor models hint at two stages of organization

To evaluate the models, we calculate the properties of obtained trajectories and assess whether they appear standard or not, if they exhibit explosive behavior (a rapid increase in the size of the largest cluster, i.e., the connected component with the largest number of nodes, denoted as C_1_), and whether they increase gradually (which would be atypical). While the trajectories for the ER, AE, TE, and LF methods follow a typical pattern of a single explosive transition, the results of other models are more complex. Due to the nature of the models, in CC and CA, there is an initial transition phase, which plateaus at some level, followed by a rapid transition that brings full connectivity. We assume this initial phase to occur when the fraction of number of nodes in the largest cluster to the total number of nodes is between *0*.*1* and *0*.*5*. For the CL model, the trajectories for the chromosomes exhibit variability. To account for these differences, we divide the chromosomes into two groups, based on the median split of the bias parameter (roughly corresponding to the time when the largest cluster size reaches *0*.*5*) - the group reaching *0*.*5* earlier, we denote as CLa (chromosome 1, 4, 6, 7, 11, 12, 15, 16, 17, 19, 20), and the one reaching this threshold later we denote as CLb (chromosome 2, 3, 5, 8, 9, 10, 13, 14, 18, 21, 22).

For CL, CC, and CA the first phase transition can be attributed to the formation and folding of smaller structures such as CCDs. This is also observed in the 3D-models which are created at various points of the trajectory. In contrast, the second phase transition’s slope is steeper and exhibits the characteristics of a first-order phase transition. This phase transition can be attributed to formation of compartments. Thus, we get two-phase transitions, an intra-compartmental providing local connectivity, and the second inter-compartmental phase transition that organizes the larger scale structure. The two compartments emerge as first and second-largest clusters that are growing along with the addition of edges, and merge as the compartments start to collapse. A characteristic thing we observe in the CL model is that the CCDs are forming only when the local loops are added. In contrast, in CA both the local loops and the distant loops are being added to the system simultaneously, so anchors distant in the genomic sequence having similar chromatin features are being brought together during the first, smooth percolation phase, during which the size of the largest cluster increases relatively slowly.

### Highly connected structures appear at different stages of the simulations

To assess quantitatively the differences in the characteristics of percolation in each model we calculate four measures (slope, critical region width, onset, and bias) for each of the chromosome-averaged percolation trajectories (see Methods), and analyze them using one-way ANOVA, followed by pairwise t-tests. We concentrate on the slope (Fig. 3B) as the primary measure of rapidness of percolation. The measures for the models with simpler trajectories (ER, AE, TE, and LF) are analyzed during the entire percolation. However, for the models with two discernable phases i.e. CC and CA, only the measures for the first phase are analyzed, denoted as CC (1) and CA(1), as the second phase is noncontinuous. For the CL model, the measures for both phases are specified separately for the two groups of chromosomes: CLa(1) denoting the first phase for the group with the initial rapid phase, while CLb(1) is the first phase for the second group of chromosomes. CLa(2) and CLb(2) represent the second phase for both groups.

**Figure 3:**
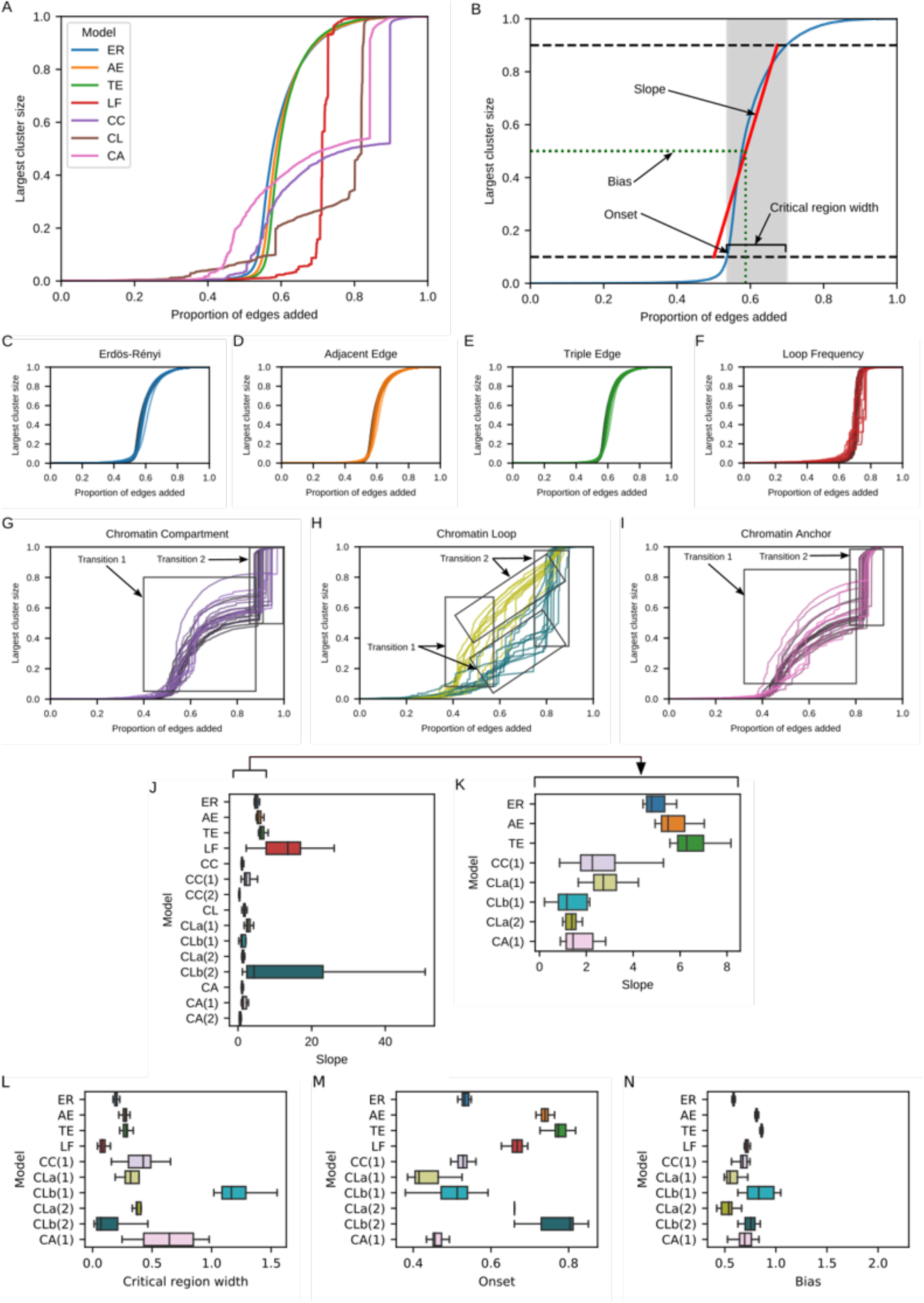
(A) The trajectories of the largest cluster size, expressed as a proportion of the total number of nodes in the graph, obtained for all the models and their statistical analysis, presented for chromosome 8 as an example. (B) A visual explanation of the measures used to quantify the trajectories. (C-I) The averaged trajectories for each chromosome for the Erdös-Rényi model (C), Adjacent Edge model (D), Triple Edge model (E), Loop Frequency model (F), Chromatin Compartment model (G), Chromatin Loop model (H), and Chromatin Anchor model (I). Each line represents an averaged trajectory of a single chromosome. (J-N) The models differ in the values of: slope (J, K), critical region width (L), onset (M), bias (N).

The results of the ANOVA omnibus test for the slope parameter are highly significant (*F(9, 166) = 13*.*15, p <* .*001*). The post-hoc tests reveal multiple significant differences between the models (all reported p-values are Bonferroni-corrected) (Supplementary Tables 1 and 2). We focus on selected interpretable differences, and the complete results can be seen in boxplot form in Fig. 3A. The LF model has a significantly higher slope than any other model (maximum *p =* .*008*, for LF-TE comparison), and for CLb(2) *(p = 1*.*0*). CLb(2), on the other hand, has larger variance and thus does not differ significantly from other models, while having similar characteristics as LF. This is indicative of many tiny separate clusters forming in the beginning, which are then rapidly merged in the critical region, in an almost-non continuous fashion. Furthermore, the three topological models (ER, AE, and TE) have increasing slopes (in that order), each pair having a statistically significant difference (*max. p =* .*008* for AE-TE). it is their expected behavior (D’Souza and Mitzenmacher, 2010) and it reinforces the confidence in the applicability of percolation analysis for our networks. These three models also have significantly higher slopes than the other models (all *p <* .*001*), except for the aforementioned LF and CLb(2). For the CC and CA models, as mentioned earlier, the general characteristics of the trajectories are similar, and the difference in slopes in the first phase is not significant (*p =* .*25*). These models in the first phase percolate slower than ER, AE and TE (as indicated by lower slopes), forming a small number of large structures that are then connected in the second phase, when a sharp second-order transition occurs. Finally, for the CL model the first phase of group “b”, i.e. CLb(1) and both phases of group “a”, i.e. CLa(1) and CLa(2) show similar behavior to the one in CC and CA models (the slopes are lower than ER, AE and TE). Additionally, as expected, the two groups of chromosomes (“a” and “b”) differ in their behavior between the two phases. The group “a” has a higher slope in the first phase than in the second phase (*p =* .*003*), while group “b” has, in contrast, a lower slope in the first phase, followed by an almost noncontinuous transition afterwards. This indicates that the structure of particular chromosomal networks differs across the chromosomes, and this difference is not readily explainable by simple properties such as size, as both groups contain both large and small chromosomes (e.g. “a” contains 1, 4 and 19, 20, while “b” contains 2, 3 and 21, 22).

The analysis of the critical region width reflects the general pattern of the result for the regression with higher slopes corresponding to more narrow critical regions (see Fig. 3B). The onset and bias (Fig. 3C-D respectively), while not directly comparable for the two-phase models (CC, CL and CA), for the topological models give results consistent with theoretical expectations (D’Souza and Mitzenmacher, 2010; D’Souza and Nagler, 2015), i.e. increasing values of onset and bias (increasingly delayed percolation) for ER, AE and TE (in that order). The differences are significant for the onset parameter (*p <* .*001*), but not significant after correction for the bias parameter, even though the order is preserved. As a final validation of our critical region calculation, we calculate the onset parameter for the trajectories of two additional system properties: the size of the second-largest cluster (C_2_), and the second raw moment of cluster sizes (m_2_), normalized to the range 0-1. In accordance with the expectations these alternative onsets indicate similar points of percolation: the Pearson correlation calculated for each model and chromosome (*N=154*) between the original onset, and the C_2_ onset is 0.81 (*p <* .*001*), and between the original onset and the m2 onset is *0*.*91 (p <* .*001)*.

To summarize, the data-driven models produce trajectories that are different from purely topological models, both quantitatively and qualitatively. These results reveal that:

1. The percolation trajectories differ between methods in terms of the rapidness of the percolation, indicating the most rapid percolation for the LF method, and replicating the patterns outlined in the theory (D’Souza and Mitzenmacher, 2010; D’Souza and Nagler, 2015)
2. CC, CL, and CA models exhibit non-uniform behavior, corresponding to different edges being added - two phases emerge related to the compartmental organization, hence by using these models we can capture the idea of genomic condensation into compartments.
3. The CL model is sensitive to chromosome structure, while others show relatively stable behavior for the different chromosomes.

### The Percolation theory with Loop Extrusion reproduces the dynamic behavior of the chromatin and the relationship between CCDs and Compartments

The dynamics of loop formation, which leads to the creation of CCDs, cannot be studied by solely using percolation models. The chromatin loops were shown to form by active extrusion by cohesin complex. Therefore, to get the full picture, we use loop extrusion along with the percolation to model the CCD. To understand the chromatin folding and the interplay between the CCDs and compartments, both the modeling for CCDs formation using loop extrusion and formation trajectory using percolation theory should take place at the same time. We propose a graph-based method using the two modeling approaches, i.e., percolation theory and loop extrusion where loop extrusion is modeled for each loop added in each step of the Percolation Model. Modeling of the loop extrusion is performed based on the idea that loops are formed in a nested manner by the cohesin movement along with the chromatin fiber (Supplementary Figure 22). The example of loop extrusion is shown in a supplementary video for chromatin region chr3:188989868-190259553. In this particular region we have 15 CTCF interactions having PET cout over four. We first used the percolation method followed by LEM to show the chromatin folding. To design the synthetic LEM data, we got 1303 CTCF motifs which were validated by Cohesin ChIP-Seq peaks. Using these CTCF motifs as the binding factor of cohesin while loop extortion. Loop extrusion represents the movement of cohesin along the chromatin fiber. Therefore, the trajectory of Loop Extrusion Model (LEM) starts to represent the real-time formation of the loop (supplementary movie). If the orientation of the CTCF motif is convergent, then cohesin stops and the anchor position is fixed; if it is divergent, it slows down. We model the dynamic loop formation and binary percolation together. At each time point of the percolation model we introduce a loop into a polymer model of chromatin 3D structure (Kadlof *et al*., 2020) and model its dynamic nature using LEM. Therefore, chromatin folding by percolation and loop extrusion allows for the 4D (3D structure + time) computational modeling of chromatin organization.

### Application of our methods to chromosome 8 of GM12878, H1ESC, and HFFc6 cell-lines

To validate our methods, we extended the current study and applied it into the latest version of the human genome i.e., hg38 for GM12878 and H1ESC, and HFFc6 cell lines. We only ran our final model i.e., chromatin anchor model over these cell lines. To obtain the scalar parameter for both anchors and loops first we ran a Monte Carlo Feature Selection (MCFS) (Dramiński and Koronacki, 2018) algorithm. This enables us to identify the important set features for anchors and nodes and use this set of features to create the scalar parameter instead of using all features. Then we studied the folding trajectories. It is shown from the trajectories for chromosome 8 for GM12878, H1ESC, and HFFc6 that the folding pattern is conserved across the cell lines.

We applied the Monte Carlo Feature Selection algorithm on the epigenetic data (for the full list see Supplementary Table 2) and identified the most important features in multi-scale classification tasks. The results are shown in **Table 1**. - Those epigenetic features were used further for the LDA. After the creation of the scalar parameter, we followed the chromatin anchor model to guide the percolation, where we used the scalar values of both nodes (anchors) and edges (loops).

**Table 1.** Most important features obtained from MCFS algorithm, used further as input to LDA (Features in bold shows common between three cell lines).

| GM12878 | H1 | HFF |
| --- | --- | --- |
| **H3K4me1** | **H3F3A** | **H3F3A** |
| **POLR2A** | **H4K8ac** | **H2AK5ac** |
| **H3F3A** | **H3K79me1** | **H2BK12ac** |
| **POLR2AphosphoS5** | H3K9ac | **H2BK15ac** |
| H3K79me2 | H4K20me1 | **H3K4ac** |
| **H3K79me1** | KDM1A | H3K9me1 |
| REST | **H2BK5ac** | H4K91ac |
| **H3K4ac** | GABPA | H3K14ac |
| BHLHE40 | DNase.seq | **H3K4me1** |
| **H3K18ac** | spin | JUND |
| **MAX** | **POLR2A** | H4K12ac |
| MAZ | USF2 | H4K5ac |
| TCF12 | **H2AK5ac** | RXRA |
| H3K9me2 | HDAC2 | H2BK5ac |
| SIN3A | H3K9me2 | H4K8ac |
| RNA.Seq.minus.total | MAX | H3K18ac |
| ELK1 | POLR2AphosphoS5 | H2BK120ac |
|  | H3K36me3 | H2BK20ac |
|  | H2BK15ac |  |
|  | H2BK12ac |  |
|  | SIN3A |  |

We used a group of nodes with similar scalar values from either set 1 or set 2. Whenever the difference between scalar parameters of the anchors was the same for two loops, we selected the one with the highest absolute value of scalar value. Having added all the loops between nodes from set 1 and set 2, we added loops associated with nodes from set 3. In CA, we guide the percolation using only the information about interactions, without strength or compartmentalisation. We observed that the critical region of the merging of two compartments is constant for all the cell lines after adding around 80% of edges, as shown in Fig 5. The results show that our methods are robust, consistent, and can capture the chromatin folding trajectories throughout various cell lines and when using features from various sources.

**Figure 4:**
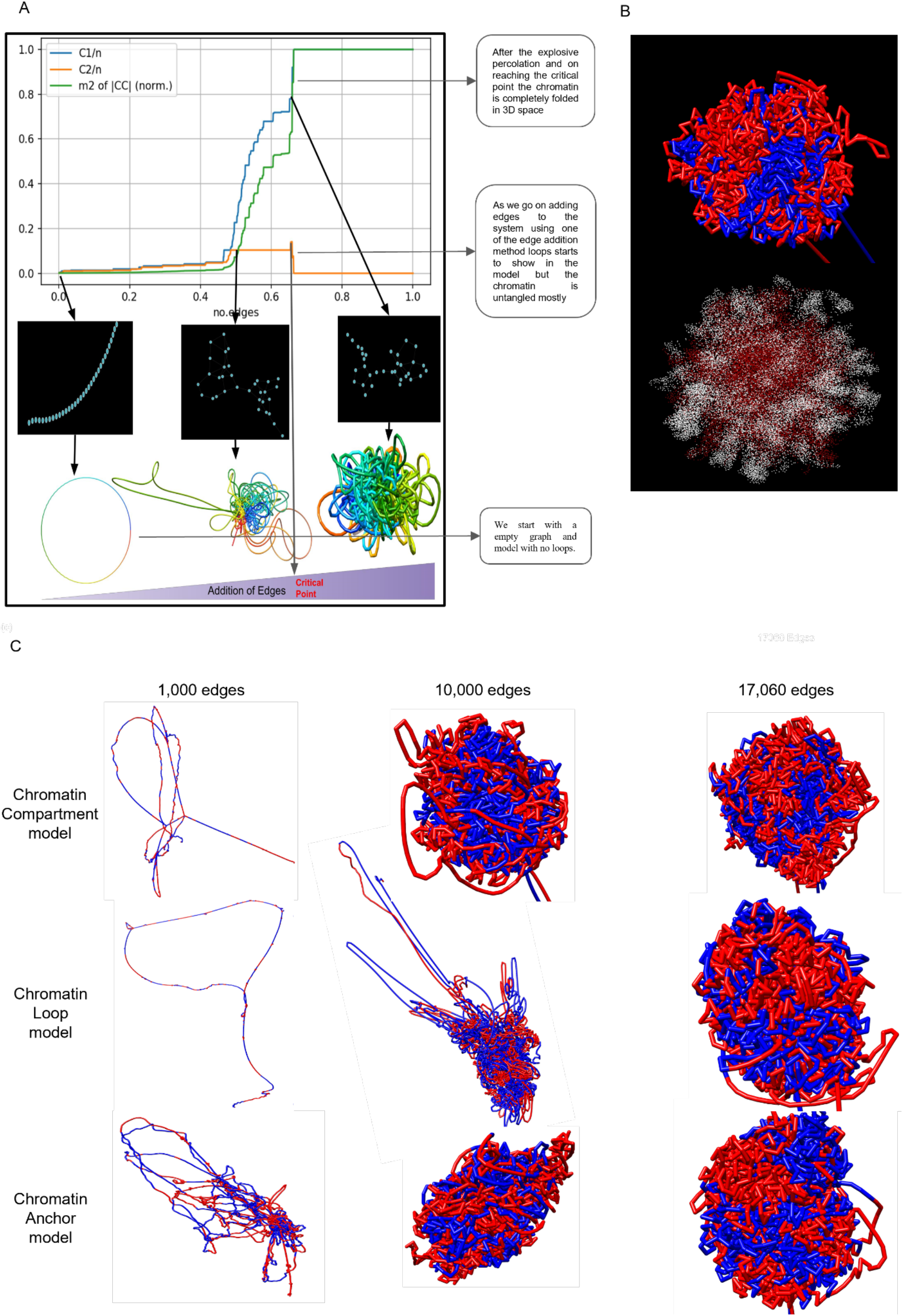
The correlation of trajectories graphs and 3D-models and how we can connect them together. (A) shows the percolation trajectory using Frequency Edge method. We start with an empty graph, so we only have a set of nodes and also the 3D model is at its initial structure with no loops. Then we continue adding edges to the network at the second point. After adding around 50% of edges we obtain a semi-connected graph, and the 3D model shows some loops, which are, however, not condensed in 3D space. Next, around the critical point we see the graph is almost connected and the 3D-model is condensed and completely folded in 3D space. (B) shows the structure of chromosome 8 (above) and the corresponding network representation of chromosome 8 using a color scheme following the compartment information from Rao et. al. In This model, compartment A is represented by red and B by blue, whereas in the graph compartment A is white and compartment B is red. (C) shows the modeling at three different time points i.e. after adding 1000, 10000, and, finally, all 17060 loops for chromosome 8 to the system for the three biological data driven models (CC, CL and CA). We can clearly see in models in CC loops that the compartment is created first. In CL we see that local loops inside a CCD are created first. In CA we can observe that the local loops as well as those loops which bring distant anchors together in 3D, due to similar epigenetic features inside CCDs, are created simultaneously.

**Figure 5:**
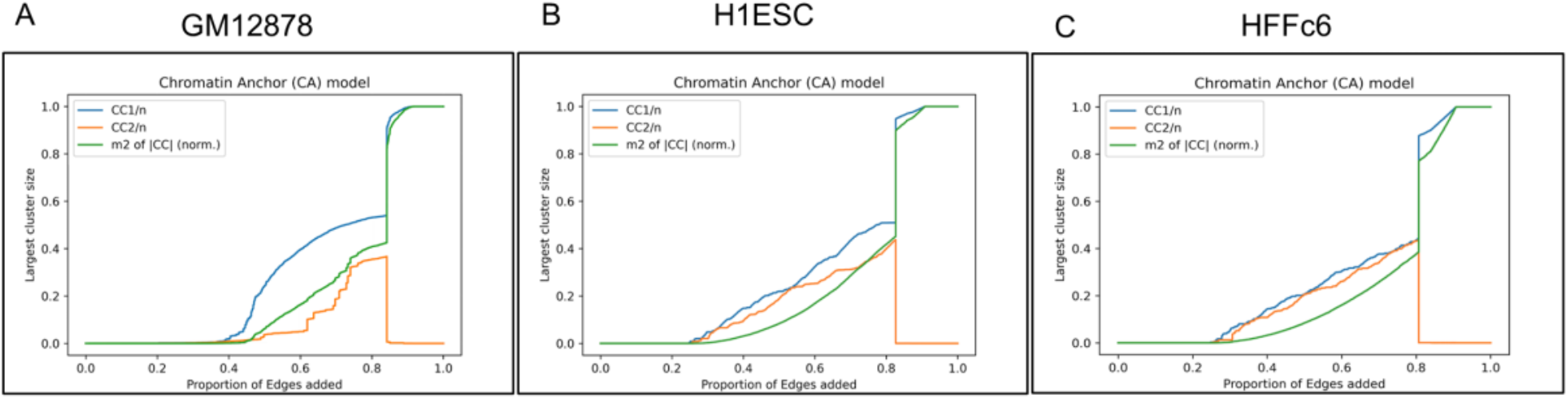
(A) The trajectories of the largest cluster size, expressed as a proportion of the total number of nodes in the graph, obtained for all the models, presented for chromosome 8 for (A) GM12878 hg38. (B) H1ESC hg38. and (C)HFFc6 hg38.

## 3. Discussion

We have proposed an application of percolation to study the formation of 3D structural organization of chromatin in the cell nucleus. Our percolation theory-based models can reproduce chromatin folding in event-based time. A graph-based approach has been introduced on CTCF mediated ChIA-PET interactions on a single-loop resolution. Moreover, we have proposed two groups of algorithms used to add edges to the graphs. The first group is based on adding edges randomly or based on the topological properties of the graphs. The methods in the second group are driven by various biological information such as PET counts of loops, compartments, genomic features, and others. We propose the order parameter - a scalar value calculated from chromatin features using Linear Discriminant Analysis, which is fitted to classify the compartments efficiently. Our approach can be applied to any genome with chromatin loop information (e.g., based on ChIA-PET or Hi-C experiments) and optional genomic information like compartments or genomic features correlated with compartmentalization. The presented models of chromosomes can be extended to the whole genome model in a straightforward manner by the addition of inter-chromosomal interactions.

To the best of our knowledge this is the first approach to applying percolation theory to chromatin structure to determine folding trajectories. We have expanded the existing models by additional biological information, reaching beyond network structure of interactions. Statistical analysis highlighted the differences and similarities between the models and revealed the more complex nature of trajectories for the data-driven models. This indicates that the large-scale network organization of the genome is coupled with chromatin features, aggregated into an order parameter, and shows a phase separation. Finally, we have successfully applied trajectories obtained from percolation models to guide the 3D modeling of the chromatin. This work can be extended to cancer cell lines to check whether they follow the same behavior in terms of chromatin folding.

## Contributions

D.P. proposed the idea of the modeling of chromatin phase separation by percolation model. K.S., M.D., A.M., R.D., and D.P. developed the initial concept further and performed the whole study. K.S., M.D., M.C., T.S., A.M., S.K. and D.P. gathered the data, implemented the algorithms, and performed the simulations, and completed the statistical analysis. K.S., T.S., M.D., M.C. and D.P. prepared the manuscript. R.D., and A.M. consulted and reviewed the manuscript. YR provided CTCF long read and in situ ChIA-PET GM12878 experimental data. All authors approved the final manuscript. K.S and M.D. contributed equally to the manuscript as co-first authors.

